# Lola-I is a developmentally regulated promoter pioneer factor

**DOI:** 10.1101/2022.04.25.489272

**Authors:** Vivekanandan Ramalingam, Xinyang Yu, Brian D. Slaughter, Jay R. Unruh, Kaelan J. Brennan, Anastasiia Onyshchenko, Jeffrey J. Lange, Malini Natarajan, Michael Buck, Julia Zeitlinger

## Abstract

While enhancers are often regulated at the level of accessibility by pioneer factors, promoters tend to be constitutively accessible and poised for activation by paused Pol II — thus are often not considered as sites of developmental regulation. Here we show that the accessibility of promoters and the acquisition of paused Pol II can be subject to developmental regulation by pioneer factors. We show that Lola-I, a *Drosophila* zinc finger transcription factor, is ubiquitously expressed at the end of embryogenesis and causes its target promoters to become accessible and acquire paused Pol II throughout the embryo. This promoter transition is required but not sufficient for tissue-specific target gene expression. Lola-I mediates this function by binding to the edges of the promoter nucleosomes, which leads to their depletion, similar to the action of pioneer factors at enhancers. These results uncover a level of regulation for promoters that is normally found at enhancers, providing further evidence that promoters and enhancers display unexpectedly similar characteristics.

## Introduction

Gene regulation during development depends on the coordinated action of enhancer and promoter sequences. Enhancers respond to specific developmental signals and transmit the information to the core promoter where transcription of the gene begins (Spitz and Furlong, 2012). Since promoters must respond to a large variety of different enhancers, they tend to have constitutively accessible chromatin (Crawford et al., 2006; Reddington et al., 2020; Thurman et al., 2012). In contrast, distally located developmental enhancers typically only become accessible in the cell lineage in which they become active (Andersson and Sandelin, 2020; Spitz and Furlong, 2012). In their natural state, developmental enhancers are inaccessible to most transcription factors, presumably to prevent activation in inappropriate cell types. The barrier arises because enhancers are often wrapped around strong nucleosomes (Adams and Workman, 1995; Simpson, 1990; Sun et al., 2015). To overcome this nucleosome barrier, so-called pioneer factors recognize their DNA-binding motifs on nucleosomal DNA and initiate chromatin remodeling (Zaret, 2020). Once enhancers are accessible, other transcription factors can bind and drive the enhancer towards activation.

Promoters on the other hand use a variety of mechanisms to maintain their accessibility across cell types and conditions. Some inherently have a low nucleosome barrier due to nucleosome-disfavoring sequences (e.g.poly-A tracts) (Field et al., 2008; Jiang and Pugh, 2009; Kaplan et al., 2009), while others are actively kept accessible through constitutively expressed pioneer factors, which antagonize the main nucleosome over the promoter. Examples for such constitutively expressed promoter pioneer factors are Reb1 or Abf1 in budding yeast, GAGA factor (GAF) in *Drosophila*, and SP1 in mammals (Biggin and Tjian, 1988; Fuda et al., 2015; Lee et al., 2007; Li et al., 1994; Okada and Hirose, 1998).

A hallmark of accessible promoters is the presence of paused RNA polymerase II (Pol II), which is present even in cell types where the genes are inactive (Ramalingam et al. 2021; Gaertner et al. 2012; Guenther et al. 2007; Muse et al. 2007; Zeitlinger et al. 2007; Gilchrist et al. 2010; Adelman and Lis 2012). At such poised promoters, Pol II initiates transcription and transcribes for 30-50 bp before going into a paused state, which promotes robust induction of genes during development (Boettiger and Levine, 2009; Lagha et al., 2013; Ramalingam et al., 2021). Upon induction, paused Pol II is released into productive elongation and new Pol II initiates at high frequencies, which results in a burst of transcription (Fukaya et al., 2016; Larsson et al., 2019; Shao and Zeitlinger, 2017).

While it may be beneficial to continually maintain promoters in a state poised for activation, the poised state of some promoters might be regulated during development. Indeed, by analyzing promoters with high levels of paused Pol II in *Drosophila*, we previously observed that a subset of promoters had no Pol II occupancy during the beginning of embryogenesis but acquired paused Pol II gradually over time (Gaertner et al., 2012). However, how such a promoter transition might be developmentally regulated and connected to gene activation was not known.

Here we now show that this group of promoters is regulated by the *Drosophila* transcription factor, Lola-I, which acts as a pioneer factor. Analogous to pioneer factors at enhancers, Lola-I functions at a step prior to gene activation. Lola-I makes its target promoters broadly accessible throughout the embryo by binding to its DNA binding motif near the nucleosome edge. This in turn leads to de novo acquisition of paused Pol II on the promoters over the course of embryogenesis, resulting in them being poised for tissue-specific gene activation during differentiation. These results illustrate a new level of regulation for promoters, one that is likely common in mammals, and provide further evidence that enhancers and promoters share some similar characteristics.

## Results

### Lola-I is required for paused Pol II and chromatin accessibility at target promoters over developmental time

We reasoned that the promoters that acquire paused Pol II at later stages of embryogenesis (defined as opening set genes in Gaertner et al., 2012) could be regulated by a transcription factor. We therefore searched for DNA-binding motifs that are enriched at these late-stage promoters compared to promoters with high levels of paused Pol II throughout embryogenesis (constant set, Gaertner et al., 2012). The most highly enriched motif was AAAGCT (> 5-fold enrichment; Supplementary table 1) (Figure 1A), a motif bound by the zinc-finger transcription factor Lola-I (Enuameh et al., 2013).

**Figure 1:**
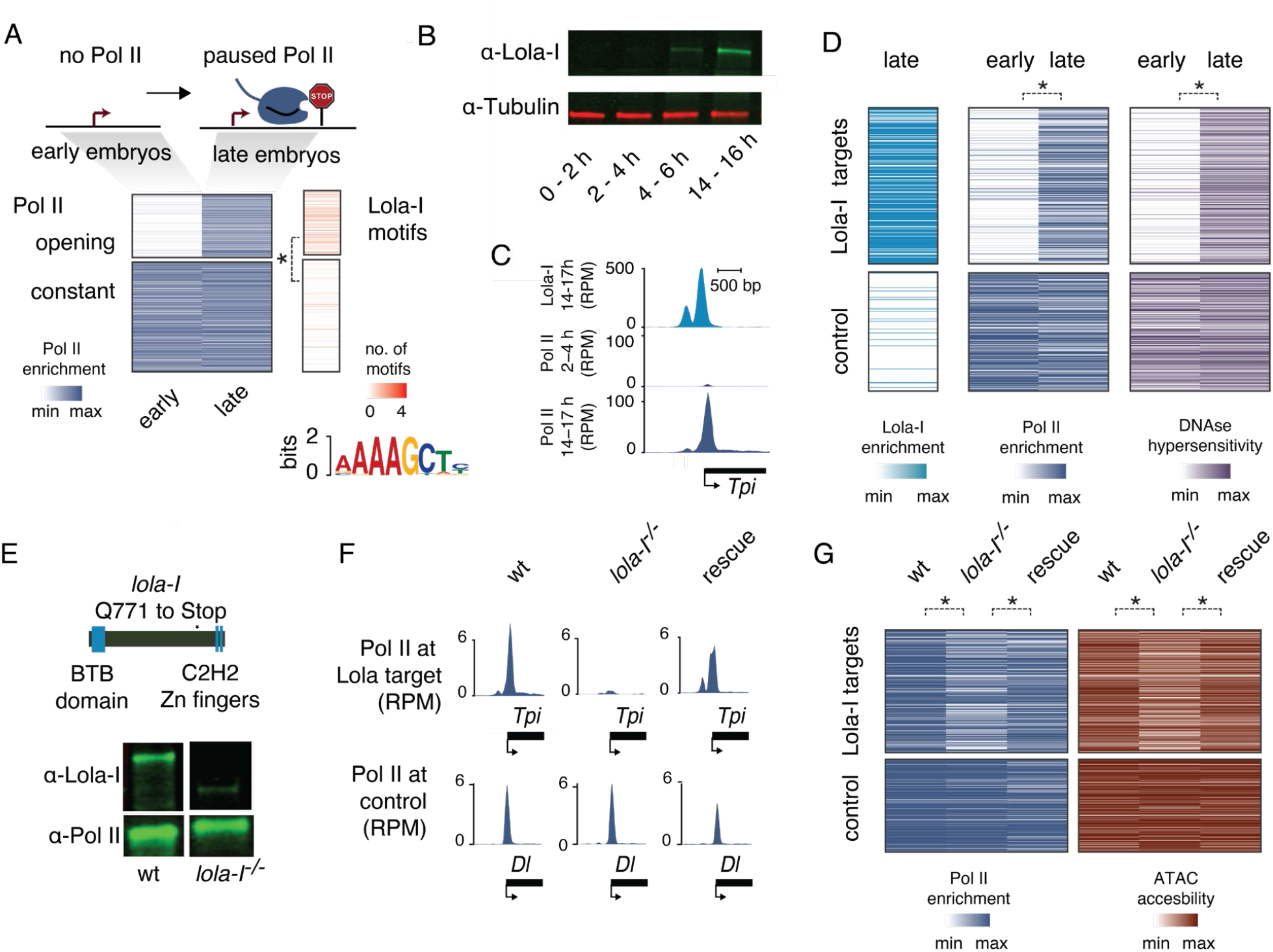
Lola-I is required for the loading of paused Pol II to target genes in the late *Drosophila* embryo. **A**) At promoters that have no Pol II occupancy in the early embryo but acquire paused Pol II in the late embryo (opening set), the Lola-I motif was identified *de novo* by MEME analysis (e-value < 1e-90). The Lola-I motif was also specifically enriched at the promoter regions (−200 bp upstream to the TSS) of the opening set versus the constant set (enrichment = 6.6, *p < 1e-33, chi-squared test with multiple testing correction). **B**) A Western blot with antibodies specific for the Lola-I isoform shows that Lola-I Increases in expression during embryogenesis. alpha-Tubulin is shown as control. **C**) Single-gene example showing that Lola-I binding (measured by ChIP-seq) is found at the promoter of *Tpi*, a gene that acquires paused Pol II over time. **D**) Heatmaps showing that genome-wide Lola-I binding at target promoters is associated with an increase in Pol II occupancy, RNA levels, and DNAse hypersensitivity from the early to the late embryo. A random sample of 250 promoters from the constant set (Gaertner et al., 2012) is used as control promoters. The star denotes significance (*P < 1e-15) using a two-sided Wilcoxon test. **E**) Mutant line *lola-I* ^*ORC4*^ *(Goeke et al., 2003)* has a premature stop codon before the C2H2 zinc-finger region that codes for the DNA binding domain. A Western blot confirms that Lola-I is a truncated product in *lola-I* ^*ORC4*^ homozygous embryos and suggests that it is degraded. The Rpb3 subunit of Pol II is shown as control below. The wt and *lola* ^*-/-*^ lanes were not run adjacently in the original gel. **F**) Pol II occupancy at the Lola-I target gene *Tpi* (but not at the control gene *Dl*) is strongly reduced in homozygous *lola*^*ORC4*^ mutant embryos. In the rescue line, which expresses *lola-I* cDNA in the *lola-I* ^*ORC4*^ mutant background, Pol II occupancy is rescued to wild-type levels. **G**) Heatmap showing that the Pol II occupancy and ATAC-seq accessibility is specifically reduced at the Lola-I target promoters in *lola-I* ^*ORC4*^ mutant embryos. In the rescue line, which expresses *lola-I* cDNA in the *lola-I* ^*ORC4*^ mutant background, Pol II occupancy and accessibility are rescued to levels comparable to wild-type. The star denotes significance (Pol II – wt-mutant: *P < 1e-5, mutant-rescue: *P < 1e-4; ATAC– wt-mutant: *P < 1e-9, mutant-rescue: *P < 1e-7) using a two-sided Wilcoxon test.

Lola-I is encoded by one of the more than 25 different splice isoforms from the *lola* locus (Goeke et al., 2003). All Lola proteins code for transcription factors that have the same N-terminal BTB domain, but also have isoform-specific C-terminal zinc-finger domains with distinct DNA-binding specificities and developmental roles (Goeke et al. 2003; Zheng and Carthew 2008; Neumüller et al. 2011; Giniger et al. 1994). Notably, the RNA of Lola-I is upregulated during the later stages of embryogenesis (Casas-Vila et al., 2017), consistent with our proposed role for Lola-I.

To test whether Lola-I indeed specifically binds to the promoters in the opening set, we raised polyclonal antibodies specific for the Lola-I isoform. These antibodies confirmed that the Lola-I protein strongly increased in abundance during the late stages of embryogenesis (Figure 1B). We then used these antibodies to perform chromatin immunoprecipitation (ChIP-seq) experiments on late-stage *Drosophila* embryos (14-17h).

We found that Lola-I bound regions are enriched for the Lola-I motif (407/500 top signal regions have a Lola-I motif within 100 bp of the peak summits, MEME motif enrichment E-value < 1e-500) and the Lola-I-bound promoters are indeed those that acquire paused Pol II over time (Figure 1C, 1D, S1A). These promoters also show increased chromatin accessibility in DNA hypersensitivity (DHS)-seq data (Figure 1D), while control promoters from the constant set (Gaertner et al., 2012) did not show this increase (Figure 1D, S1A). We also observed specific conservation of Lola-I binding motifs across *Drosophila* species, which is indicative of their functional importance (Figure S1A, B).

To test whether Lola-I is the factor that causes these promoters to change over developmental time, we analyzed a previously identified *lola-I* mutant, *lola*^*ORC4*^ (Goeke et al., 2003). This mutant contains a premature stop codon specifically in the *lola-I* isoform, leading to a truncated Lola-I protein without the zinc-finger DNA-binding domain (Figure 1E), which we confirmed by Western blot (Figure 1E). We found that in these *lola-I* ^-/-^ embryos, Pol II occupancy, as measured by ChIP-seq experiments, was specifically reduced at Lola-I targets, but not at control promoters (Figure 1F, 1G). Furthermore, Lola-I targets showed reduced chromatin accessibility as measured by ATAC-seq (Figure 1G). These defects were not due to a developmental delay or due to secondary mutations in the *lola* ^*ORC4*^ line (Figure S1C) and were rescued by the transgenic expression of Lola-I (Figure 1F, 1G). This demonstrates that Lola-I mediates the acquisition of paused Pol II and chromatin accessibility specifically at the promoters to which it binds.

### Lola-I establishes paused Pol II throughout the embryo but mediates tissue-specific gene expression

The simplest explanation for the observed effect of Lola-I on Pol II is that Lola-I is a strong activator that opens promoters and leads to increased levels of Pol II recruitment and productive elongation. If this were the case, binding of Lola-I would be expected to correlate both temporally and spatially with the expression of its target genes. On the other hand, occupancy of Lola-I and paused Pol II in tissues where the target genes are not expressed would argue that Lola-I establishes paused Pol II independently of gene activation.

To distinguish between the two scenarios, we first analyzed where Lola-I is expressed in the embryo. In immunostainings, nuclear Lola-I was found ubiquitously throughout the embryo (Figure 2A, S2A). We also detected ubiquitous *lola-I* RNA by single-molecule RNA fluorescence *in situ* hybridization (single-molecule FISH) (Figure S2B). With Lola-I being ubiquitous, we next asked whether paused Pol II was also acquired ubiquitously across all tissues. To isolate specific tissues from late-stage embryos, we used the INTACT method (Bonn et al., 2012; Deal and Henikoff, 2011; Ramalingam et al., 2021) and performed Lola-I and Pol II ChIP-seq experiments on the isolated nuclei.

**Figure 2:**
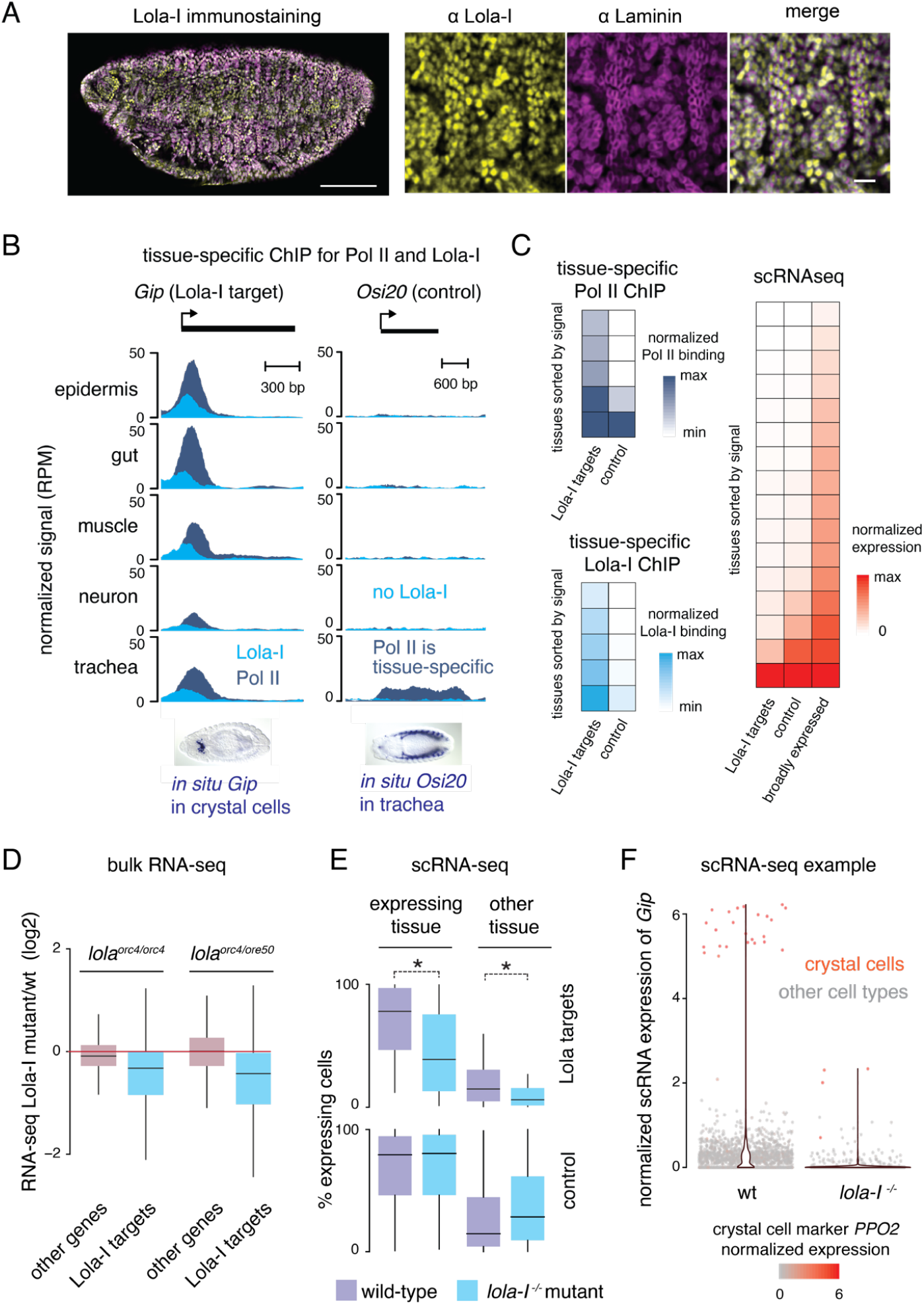
Lola-I establishes paused Pol II throughout the embryo and promotes robust tissue-specific gene expression. **A**) Immunostaining using the Lola-I antibodies (yellow) shows that Lola-I is expressed ubiquitously in the late *Drosophila* embryo. Ubiquitous Lamin is shown as control (pink), which also shows the nuclear localization of Lola-I. Here the brightness and contrast settings of the linear look up table are linearly adjusted for clarity. The settings in the individual panels are the same as in the merge. **B**) ChIP-seq experiments on isolated embryonic tissues using either Lola-I antibodies (turquoise) or Pol II antibodies (dark blue) reveals that Lola-I binding and paused Pol II are found in all examined tissues, even when the gene is expressed in a specific tissue only (*Gip* is shown as example). Ubiquitous Pol II is not found for all genes, e.g., the control gene *Osi20* shows Pol II binding and expression only in tracheal cells. ChIP-seq data are shown as normalized reads per million (RPM). The tissue-specific expression of *Gip* in crystal cells and *Osi20* in tracheal cells are known from *in situ* hybridization shown below (courtesy of Berkeley *Drosophila* Genome Project (Tomancak et al., 2002, 2007)). **C**) Average Pol II occupancy, Lola-I occupancy, and scRNA-seq levels confirm that Lola-I target genes show paused Pol II broadly across tissues but show tissue-specific gene expression, similar to the expression of the previously described control genes with the most restricted Pol II occupancy across tissues (Ramalingam et al. 2021). The Pol II signals are normalized to the maximum value across tissues for each gene and tissues are sorted from low to high average signal. Normalized mean profile for each group is obtained by averaging the sorted normalized profiles of all genes in a group. Similarly, expression values are normalized across tissues. Lola-I binding signal is not normalized across tissues for each gene to enable comparison of Lola-I binding between the Lola-I target genes and other tissue-specific promoters. **D**) Boxplot of bulk RNA-seq data show that Lola-I targets are down-regulated in *lola-I* mutants compared to wild-type (14-17 h), while all other expressed genes show overall similar expression levels. This confirms that Lola-I mediates its effect by directly binding to target promoters, and not by indirect effects that would affect all genes (Wilcoxon two-sided test, *P<1e-10). **E**) Boxplots of scRNA-seq data show that both the tissue-specific expression and the basal expression of Lola-I targets are reduced in the *lola-I* embryos compared to wild-type embryos (both profiled at 14-14.5 h) (Wilcoxon two-sided test, expressing tissue: *P<1e-4, other tissue: *P<1e-2). Expression is shown as the fraction of cells with detectable expression. The same trend is observed for the median expression (Figure S1G). **F**) scRNAseq expression data for the Lola-I target gene *Gip* is shown for each cell (grey dots) isolated from wild-type or *lola-I* mutant embryos. A violin plot of the data (red) is laid on top. In wild-type embryos, *Gip* is expressed in the small number of crystal cells (marked by the presence of the *PPO2* gene) at high levels, but also shows background expression in the majority of cells, where *Gip* is not expected to be expressed. In *lola-I* embryos, both the specific expression and the basal expression are reduced. Box plots in D and E show the median as the central line, the first and the third quartiles as the box, and the upper and lower whiskers extend from the quartile box to the largest/smallest value within 1.5 times of the interquartile range.

Lola-I binding and paused Pol II were present at target promoters across all examined tissues. This was true even for tissue-specific genes (Figure 2B, 2C). For example, the gene *Gip* is specifically expressed in crystal cells, an immune cell type found near the proventriculus, as observed by *in situ* hybridization of whole-mount embryos. However, paused Pol II is present at this locus in all tissues, not just in crystal cells (Figure 2B). This is not due to heterogeneity or low purity of isolated tissues since control genes show the expected tissue specificity of Pol II (*Osi20* in Figure 2B) (Ramalingam et al., 2021). This suggests that Lola-I changes the promoter state without necessarily activating gene expression.

To confirm the ubiquitous effect of Lola-I on promoters and the tissue-specific target gene expression, we analyzed Lola-I ChIP-seq and Pol II ChIP-seq data more globally and compared the data to single-cell RNA-seq data (scRNA-seq) from an equivalent stage. As expected, Lola-I target genes showed Pol II occupancy very broadly among cell types, as is known for paused genes (Ramalingam et al., 2021). These promoters also had high chromatin accessibility (Ramalingam et al., 2021), hich was dependent on Lola-I in each of the specific tissues we isolated from *lola-I* ^-/-^ embryos (Figure S1D). In contrast, the expression of the target genes was highly tissue-specific, similar to genes with tissue-specific elongating Pol II (Ramalingam et al., 2021) (Figure 2C). This suggests that Lola-I establishes paused Pol II at target promoters throughout the embryo but that the tissue-specific expression of the genes is acquired separately, ruling out the possibility that Lola-I is simply a strong activator.

Since Lola-I binding is not sufficient to induce the expression of target genes, we next asked whether Lola-I is nevertheless required for their expression. This would imply that Lola-I-mediated chromatin accessibility and paused Pol II at promoters are a necessary step towards gene activation. By performing bulk RNA-seq experiments on *lola-I* ^-/-^ and wild-type embryos, we found that the expression of Lola-I target genes, but not that of control genes, was significantly reduced in *lola-I* ^*-/-*^ embryos (Figure 2D). These target genes were enriched for genes involved in metabolism and ion transport (Supplementary table 2). This is consistent with the mutant phenotype of *lola-I* ^*-/-*^ embryos, which fail to hatch at the end of embryogenesis but lack any visible gross abnormalities in neuronal, muscle and glial structures (Figure S2C).

Since the Lola-I target gene expression was reduced in *lola-I* ^*-/-*^ embryos, we wondered whether this loss specifically stems from the tissue where the gene is normally expressed at a high level, or whether the transcript loss could also come from changes in basal expression in other tissues. This is plausible since inactive promoters with paused Pol II typically have detectable basal transcript levels (Ramalingam et al. 2021; Gaertner et al. 2012; Lis 1998; Gilchrist et al. 2012). We performed scRNA-seq experiments and found that indeed both the tissue-specific expression and the basal levels of target genes were reduced in *lola-I* ^*-/-*^ versus wild-type embryos (Figure 2E, S1E, S1F, S1G). For example, *Gip* ‘s high expression in crystal cells and its basal expression in other tissues were both reduced in *lola-I* ^*-/-*^ mutants (Figure 2F). Within the limits by which low expression of genes can be confidently compared between two scRNA-seq samples, these results suggest that Lola-I has a general effect on promoters, impacting basal activities of promoters and enhancing tissue-specific gene induction.

### Lola-I ‘s effect on promoters increases the gene activation frequency

If Lola-I is not a traditional activator and primarily affects the promoter state, we wondered how Lola-I regulates the transcription of its target genes. The transcription of most genes occurs in bursts, characterized by a model with alternating periods of transcriptional inactivity (OFF state) and activity (ON state), during which Pol II produces a burst of multiple nascent RNAs (with a rate constant K_prod_). Lola-I could regulate the transition to the ON state (through the activation rate constant K_on_), and thereby affect the burst frequency, and/or Lola-I could regulate the burst size, which depends on the rate constants K_prod_ and K_off_ (Pichon et al. 2018; Munsky et al. 2012; Bartman et al. 2016; Zoller et al. 2018). The activation rate and burst frequency can increase as a result of enhancer-mediated gene activation (Nicolas et al. 2018; Nicolas et al. 2017; Fukaya et al. 2016; Zoller et al. 2018; Keller et al. 2020). The burst size may also increase with higher expression (Dar et al. 2012; Zoller et al. 2018; Berrocal et al. 2020) but it is also a promoter-specific property (Carey et al., 2013; Pimmett et al., 2021; Yokoshi et al., 2022). If Lola-I affects the burst size, we would expect that most cells in *lola-I* ^-/-^ embryos show a similar proportional reduction of Lola-I target gene expression. Such a result could also be observed if the gene is transcribed continuously with a certain probability, rather than in bursts (Dar et al., 2012). On the other hand, if Lola-I affects the burst frequency, we would expect a more heterogeneous reduction of transcripts in *lola-I* ^-/-^ embryos, where some cells in the mutant embryos show high levels of RNA similar to wild-type embryos, while others show very few or no RNA.

To test this, we performed single-molecule FISH (Femino et al., 1998; Raj et al., 2006) using a series of fluorophore-labeled small probes against the Lola-I target gene *Gip*. As a control, we used probes against *PPO1* and *PPO2*, which are not Lola-I target genes but are also specifically expressed in crystal cells. We found that in wild-type late-stage embryos (12-14h), *Gip* was expressed at high levels and localized to the same cells as *PPO1*/*PPO2* (Figure 3A, 3B, S3A), confirming expression in crystal cells. We then estimated the number of *Gip* RNAs for each cell by measuring the cell ‘s total fluorescence intensity and dividing it by the average intensity of individually measured RNAs (example in Figure 3A inset), which yielded an average of 720 *Gip* RNAs per cell. We also observed clearly detectable bright spots at the sites of nascent transcription (Figure 3A inset). These spots were present in 41% of the crystal cells and were not observed in other cell types.

**Figure 3:**
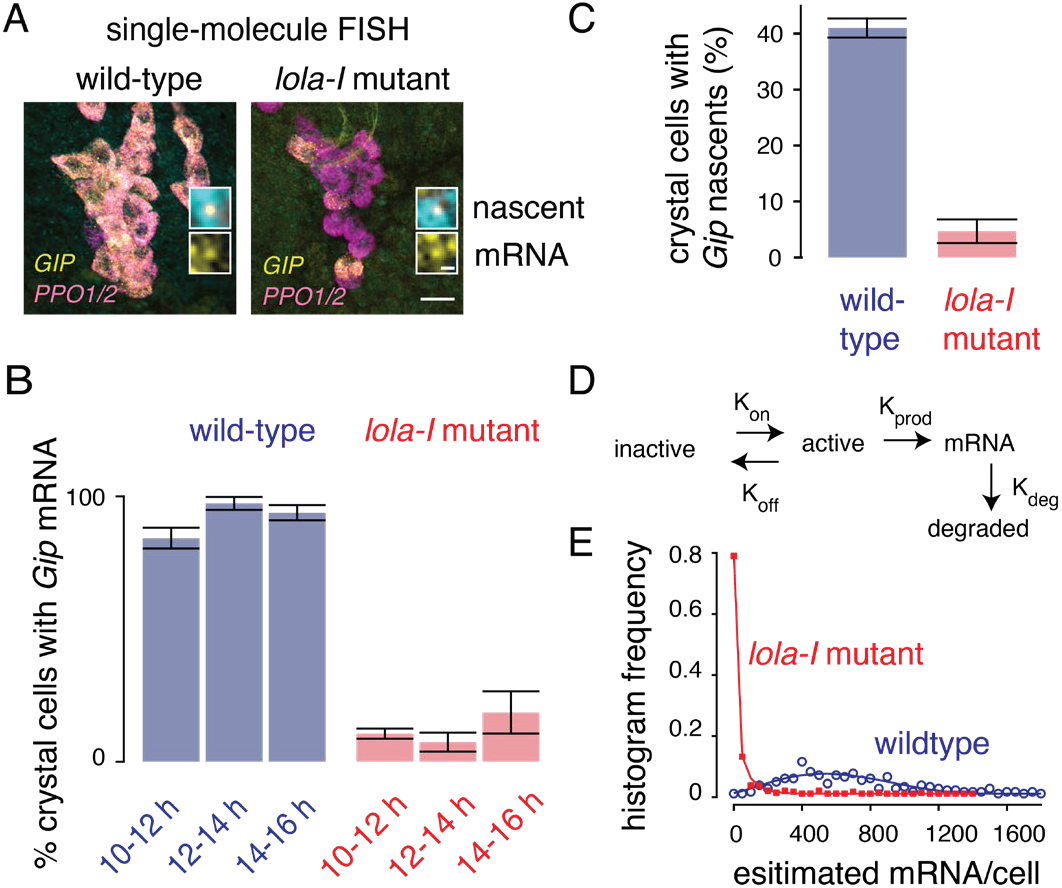
Single-molecule FISH reveals a higher gene activation barrier in *lola-I* mutants. **A**) Example of single-molecule FISH images of *Gip* (yellow), and *PPO1* and *PPO2* genes (pink) in wild-type embryos and *lola-I* ^*-/-*^ mutant embryos. Data was acquired with 40x magnification; scale bar - 20 μm. Top insets show the nascent *Gip* signal in the nuclei. Bottom insets show the single *Gip* RNAs in the cytoplasm. Here the brightness and contrast of the linear lookup table was adjusted linearly for clarity. In the overview images, the brightness and contrast settings are the same. These settings in the insets, are adjusted individually for clarity. **B**) The percentage of crystal cells (defined by high expression of *PPO1* and *PPO2*) that express high levels of *Gip* is strongly reduced in lola-I mutant embryos compared to wild-type. Barplots show data for wild-type: 10-12 h n=7, 12-14 h n=8, 14-16 h n=4, *lola-I* mutant: 10-12 h n=7, 12-14 h n=7, 14-16 h n=7, where n is the number of embryos, with the bar representing mean and the whiskers showing the standard deviation. Reduced *Gip* expression is observed for multiple time points showing that it is not due to a developmental delay and that *Gip* expression only mildly recovers over time. **C**) The percentage of *PPO1* and *PPO2* positive crystal cells with a visible *Gip* nascent site of transcription (indicative of a transcriptional burst) is also strongly reduced in *lola-I* mutants (12-14 h). The bar represents the mean, and the whiskers show the standard deviation (wild-type n=7 and *lola-I* mutant n=14, where n is the number of embryos. **D**) Two-state model that was fitted to the data. **E**) Histograms of mRNA/cell (dots), calculated from fluorescent intensities at 100x magnification (see Methods), and fitted lines. The ratio of Koff and Kon was fixed to the ratio of crystal cells with visible nascent transcripts between wild-type and mutants as shown in C.

In *lola-I* ^*-/-*^ embryos of the same stage and analyzed in the same way as wild-type embryos, *Gip* expression was notably reduced, but the reduction was not uniform across all crystal cells. While most *PPO1*/*PPO2*-positive crystal cells showed little to no detectable *Gip* expression, typically a few cells still showed strong *Gip* expression, albeit lower than wild-type levels (average of top 10% = 309 RNAs) (Figure 3A, 3B, S3A). Consistent with this, only 3.5% of the *PPO1*/*PPO2* positive cells showed bright spots of nascent *Gip* transcription in *lola-I* ^*-/-*^ embryos, compared to 41% in wild-type (Figure 3C). Hence, much fewer mutant crystal cells were actively transcribing *Gip*, but those that were had a substantial number of transcripts. To rule out that the reduced *Gip* expression was due to developmental delays, we performed the same single-molecule FISH experiments in wild-type and *lola-I* ^*-/-*^ embryos over several time points (10-12 h, 12-14 h, 14-16 h). This confirmed the reduced *Gip* expression in *lola-I* ^*-/-*^ embryos across all time points (Figure 3B, S3B).

The heterogeneous reduction in *Gip* expression across cells could be explained by a reduced burst frequency. To test this, we fitted our data from wild-type and *lola-I* ^-/-^ embryos to a simple two-state model (Peccoud and Ycart 1995), using a mathematical framework to fit the parameters K_on_, K_off_ and K_prod_ to steady-state transcript measurements with a fixed RNA degradation rate (Raj et al., 2006) (Figure 3D). Since we have measurements for the fraction of cells in the ON state from the nascent transcription spots, we fixed the ratio between the state transition rates (K_on_ and K_off_) to the ratio of cells with and without nascent transcripts. The two-state model produced a reasonably good fit (Figure 3E, chi squared goodness of fit for wild-type: 4.46 and for mutant: 3.59), and thus did not justify the added complexity of a three-state model (Bartman et al., 2019; Pimmett et al., 2021). We note that this does not eliminate the possibility of a three-state model – it simply indicates that the two-state model is sufficient to adequately fit the data.

The most striking difference between the models from wild-type and lola-I ^-/-^ embryos was the activation rate constant K_on_, which together with K_off_ determines the burst frequency (Zoller et al. 2018). We observed in the mutants a 13.8-fold reduction of K_on_ (burst frequency 8.5 fold), a 1.4 fold increase in K_off_, and a 2.7-fold reduction in K_prod_ (burst size 3.7 fold). Thus, our data also suggest a reduction in the burst size, which is consistent with previous observations that the burst size moderately decreases with lower transcriptional levels (Dar et al. 2012; Zoller et al. 2018; Berrocal et al. 2020). However, we note that very few cells are producing detectable levels of *Gip* in the mutant, thus it is difficult to accurately assess K_prod_ or burst size (Figure S3D). A live imaging system (such as the MS2-MCP) could provide a better estimate of these values. In summary, we conclude that Lola-I strongly affects the activation rate and burst frequency at its target genes. This is interesting since Lola-I acts on promoters and is insufficient for gene activation on its own. We therefore propose that Lola-I potentiates enhancer-mediated gene activation.

### Lola-I is a developmentally regulated promoter pioneer factor

To understand the mechanism by which Lola-I establishes paused Pol II at promoters, we considered whether Lola-I is a pioneer factor, which implies that Lola-I removes nucleosomes. By removing the promoter nucleosome, Lola-I would increase chromatin accessibility and allow Pol II recruitment and pausing. Pioneer factors such as GAGA factor have been shown to have such a role at constitutively open promoters (Fuda et al., 2015; Okada and Hirose, 1998), but such a mechanism has not been described for developmentally regulated promoters. To test this idea, we performed MNase experiments on early and late *Drosophila* embryos in both wild-type and *lola-I* mutant embryos. We found that Lola-I target promoters showed high nucleosome occupancy in the early embryo and a decrease in the late embryo that was Lola-I-dependent (Figure 4A).

**Figure 4:**
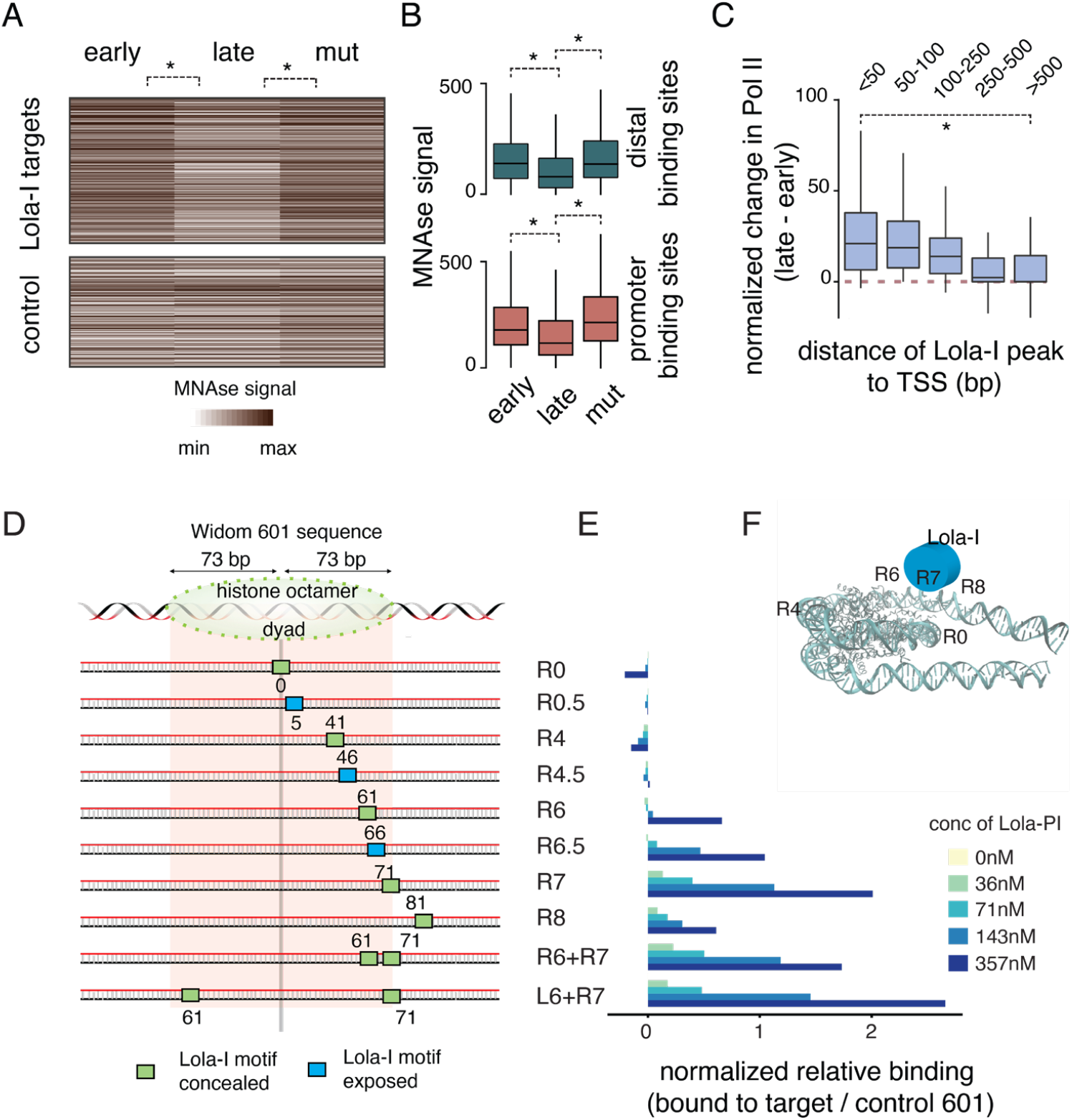
Lola-I is a promoter pioneer factor. **A**) Heatmap of normalized MNase data shows specific loss of nucleosome occupancy at the Lola-I target promoters during the late stages of *Drosophila* embryogenesis (Wilcoxon two-sided test, *P < 1e-8). **B**) Box plot shows the MNase signal centered on the Lola-I peaks from early wild-type embryos and late wild-type and *lola-I* mutant embryos. There is a decrease in nucleosome occupancy at both promoter-proximal regions and distal Lola-I binding sites in the late-stage wild-type embryos. Box plot shows the median as the central line, the first and the third quartiles as the box, and the upper and lower whiskers extend from the quartile box to the largest/smallest value within 1.5 times of the interquartile range (Wilcoxon two-sided test, *P < 1e-7). **C)** Pol II recruitment, shown as change in normalized Pol II enrichment from the early to the late stage, only occurs when the Lola-I peak is found in close proximity to the TSS, showing that Lola-I depletes nucleosomes independently of Pol II recruitment (Wilcoxon two-sided test, *P < 1e-8). **D**) Lola-I binds its motif preferentially at the edge of a nucleosome *in vitro*. Purified full-length Lola-I protein expressed in insect cells by baculovirus is incubated *in vitro* at different concentrations with nucleosomes reconstituted with Lola-I binding sites embedded in the 601 Widom sequence at different positions. **E)** The relative binding of the Lola-I protein to the 601-templates containing the Lola-I motif vs the control 601-template without the motif is measured as the loss of signal in the nucleosomal band or the gain of signal in the super-shifted band (Figure S4) (relative to the no Lola-I lane - see methods). The results show that Lola-I binds most strongly when the motif is located along the nucleosomal edge. Significant cooperativity with additional motifs is not observed. **F**) Illustration of the preferred position of the Lola-I motif with regard to the nucleosome structure.

While these results support our hypothesis of Lola-I as a pioneer factor, it is also possible that Lola-I primarily functions to recruit Pol II, and that the nucleosome depletion is a secondary effect of paused Pol II keeping the nucleosome away (Gilchrist et al., 2010; Hsu et al., 2015). To distinguish between these two scenarios, we took advantage of the fact that approximately 40% of Lola-I-bound regions are located distally to annotated promoters. These regions also contain conserved Lola-I motifs (Figure S1A, S1B), suggesting that they are functional. If nucleosomes can be removed by Lola-I, they should also be removed in these distal regions without the help of paused Pol II. Indeed, at both promoters and distal regions, Lola-I binding was associated with strong depletion of nucleosomes (Figure 4B). Nevertheless, paused Pol II was only detected at promoters, with decreasing Pol II levels the further the distance to the nearest transcription start site (TSS) (Figure 4C and S4A). This suggests that Lola-I primarily serves to deplete nucleosomes and that the recruitment of Pol II is a secondary step at promoters.

By definition, pioneer factors are able to access their motifs in nucleosomal DNA. They may do so by binding DNA at a particular position on the nucleosome, to linker DNA between nucleosomes, or at the edge of nucleosomes to DNA that becomes accessible when nucleosomes spontaneously unwrap (Li and Widom, 2004; Sekiya et al., 2009; Zaret, 2020; Zhu et al., 2018). We therefore asked whether Lola-I motifs have a preferred position on nucleosomes. Using the MNAse-seq data from early embryos where the promoter nucleosomes are still present, we found a trend of Lola-I motifs to be found at the edge of nucleosomes (Figure S4B).

Based on these results, we set out to analyze the binding preference of purified Lola-I protein to nucleosome-bound DNA using a traditional *in vitro* binding assay combined with high-throughput sequencing (Yu and Buck, 2019). Multiple DNA variants, each with a distinct position of the Lola-I motif embedded in a strong nucleosome-positioning sequence (Widom 601), were reconstituted with nucleosomes *in vitro* and incubated with different concentrations of full-length Lola-I protein (Figure 4D, 4E). The bound and unbound nucleosomal fractions were then sequenced.

The results show that Lola-I bound strongest when the motif was located near the nucleosome edge (R6, R6.5, R7 in Figure 4D, 4E, S4D, S4E) and weakest when the motif was found near the dyad (R0, R0.5, R4, R4.5 in Figure 4D, 4E, S4D, S4E). We did not find strong synergistic effects when multiple Lola-I sites were present (Figure 4D, 4E, S4D, S4E). Moreover, binding to the nucleosome edge occurred whether or not the Lola-I motif was facing outside to the solvent side or was predicted to be concealed inside (Figure 4D, 4E, S4D, S4E), suggesting that Lola-I binds when the DNA is partially unwrapped. Interestingly though, the variant with the Lola-I motif near the accessible linker region (R8) was not bound the highest, indicating that Lola-I does not necessarily prefer free DNA but may also interact with the nucleosome. These results suggest that Lola-I can best access its motif on nucleosomes when located near the edge (Figure 4F), consistent with our observations *in vivo*. Taken together, the results strongly support the conclusion that Lola-I is a developmentally regulated promoter pioneer factor.

## Discussion

In this study, we set out to identify the mechanism by which a set of promoters (Gaertner et al., 2012) acquires paused Pol II de novo over the course of embryonic development. Promoters are generally considered to be constitutively open and to have paused Pol II. Hence, we wondered what regulatory step controls the acquisition of Pol II pausing and how this change relates to gene activity. We found that the promoters are targeted by the zinc-finger transcription factor Lola-I, which is ubiquitously induced throughout the late embryo, but that the target genes are only induced in specific tissues. By facilitating the acquisition of paused Pol II, Lola-I acts at a step prior to gene activation: Lola-I ‘s function is required but not sufficient for target gene expression.

Our study therefore shows that Pol II transcription can depend on multiple limiting steps at some promoters. This is important since it is often debated whether Pol II initiation or pause release are the limiting regulatory steps for gene expression, and this is difficult to distinguish as they depend on each other (Gressel et al., 2019; Shao and Zeitlinger, 2017). Here we found that Lola-I promotes Pol II initiation and pausing at inactive promoters, which is a different regulatory step from gene activation, during which Pol II transcribes in rapid succession as part of a burst. This distinction is clear because the first step affects basal transcription and occurs ubiquitously in the entire embryo, while the second step impacts tissue-specific expression, presumably through tissue-specific enhancers that are in the promoter-proximal region or located further distally.

If Lola-I is required but not sufficient for tissue-specific gene activation, then what is its role? Our single-molecule FISH analysis suggests that Lola-I increases the rate of gene activation, which we interpret as increased responsiveness to tissue-specific activation signals from enhancers. It is tempting to speculate that this increased responsiveness is because of paused Pol II. Paused Pol II is associated with more synchronous and robust gene activation (Boettiger and Levine, 2009; Lagha et al., 2013; Ramalingam et al., 2021), with higher interaction frequencies with enhancers (Ghavi-Helm et al., 2014) and localization to the active compartment (Gu et al., 2021). Furthermore, the Downstream Promoter Element (DPE) sequence, a hallmark of promoters with Pol II pausing, increases the burst frequency (Hendrix et al., 2008; Yokoshi et al., 2022). However, Pol II pausing is highly correlated with promoter accessibility, thus we cannot rule out that some of these properties are at least in part due to increased chromatin accessibility. For example, budding yeast does not have Pol II pausing, and yet promoters that have nucleosome-repelling sequences or motifs for pioneer factors have increased promoter strength (Kubik et al., 2017; Sharon et al., 2014). These results suggest that paused Pol II and promoter accessibility could both increase the rate of gene activation.

The importance of promoter accessibility became apparent when we analyzed the mechanism by which Lola-I establishes paused Pol II. Rather than recruiting Pol II, our data suggest that Lola-I functions at promoters primarily as a pioneer factor that depletes nucleosomes. This idea is supported by our observation that Lola-I also depletes nucleosomes at distal regions where paused Pol II is not detected. Furthermore, incubation of Lola-I protein with nucleosome-bound sequences in vitro suggested that Lola-I preferentially binds to its motifs near the nucleosome edge when the DNA is partially unwrapped. Although there is no consensus on whether pioneer factors are defined by a certain binding mode in vitro (Klemm et al., 2019), such binding preference has previously been observed for some pioneer factors (Yu and Buck, 2019; Yu et al., 2021; Zhu et al. 2018). Like for other pioneer factors, binding of Lola-I may lead to the recruitment of chromatin remodelers that remove the nucleosome (Judd et al., 2021; Zaret, 2020).

Lola-I ‘s regulation of promoters at the level of accessibility is reminiscent of the regulation of enhancers. Enhancers also become accessible in a broader developmental context and through a series of regulatory events become active in very tissue-specific patterns. Enhancers are also by themselves not sufficient for gene expression, but “enhance” the expression of a gene from a promoter. Thus, Lola-I target promoters function in some way like enhancers, except that they are directly located at core promoters that facilitate the initiation and strong pausing of Pol II.

These results further blur the distinction between enhancers and promoters. The traditional separation between enhancers and promoters has already been challenged in recent years since their function is often not cleanly separable in reporter assays (Andersson and Sandelin, 2020; Haberle and Stark, 2018). However, the constitutive accessibility of promoters and the more dynamic accessibility of enhancers have still been considered to be distinctive features of these classes of elements (Andersson and Sandelin, 2020). Our finding that the accessibility of promoters can also be dynamically regulated by pioneer factors during development further supports the idea that enhancers and promoters share some fundamentally similar characteristics with each other.

Due to the general perception of promoters being constitutively open, regulation of promoters by pioneer factors is likely an understudied regulatory step. The *lola* locus has many isoforms that have different sequence-specific DNA binding domains and are expressed in various patterns during development (Goeke et al., 2003). Some of them have recently been suggested to regulate chromatin accessibility in the *Drosophila* brain (Janssens et al., 2022). By analogy to Lola-I, it is conceivable that other isoforms of Lola also target promoters and help establish paused Pol II in specific tissues or stages of development.

Promoter pioneer factors are likely to participate in regulating gene expression in other species, including mammalian development. We likely discovered this feature in *Drosophila* because the gene structure is simpler, promoters have high levels of Pol II pausing, and time course experiments can be easily performed. However, the strength of promoters in mammals can be also modulated by motifs of transcription factors (Nguyen et al., 2016). Furthermore, an opening of promoters such that they are ready for activation has been observed during early mouse development (Lu et al., 2016). An even more intriguing possibility is that promoter pioneer factors regulate the usage of alternative start sites. At least 15% of protein-coding genes in the human genome use alternative tissue-specific promoters that are enriched for specific transcription factor binding motifs (Demircioğlu et al., 2019), but the mechanism underlying this promoter selection is not known (Haberle and Stark, 2018). It is tempting to speculate that promoter selection during mammalian development is also regulated by stage-specific and tissue-specific promoter pioneer factors with analogous roles to Lola-I.

## Materials and Methods

### Fly stocks and genetic crosses

*Oregon-R* was used as the wild-type strain. Lola-I mutant lines were obtained from Bloomington stock center (ORC4 - 28267) and from Edward Giniger (ORE50). Homozygous *lola-I* mutant files were non-viable and were maintained over a *CyO-GFP* balancer to allow sorting of the homozygous mutant embryos that are GFP^-^. Lola-I rescue lines were generated as follows: a construct with an *Actin* promoter driving full-length *lola-I* cDNA and marked by *mini-white* was integrated into the *attP40* locus on 2L and then crossed with the *lola-I* ^*ORC4*^ line (*lola* is on 2R) to obtain females that recombine the second chromosome in the germ line. After crossing in a *CyO* balancer, several males harboring the *mini-white* marker were selected. After mating single males with a *CyO* balancer stock, each male was screened for the *lola-I* ^*ORC4*^ mutation by amplifying the relevant portion of the *lola* locus by PCR and sequencing. Meiotic recombinants that had both the rescue construct and the *lola-I* ^*ORC4*^ mutation were viable as homozygotes. For the INTACT experiments, embryos from fly stocks expressing tissue-specific *RAN-GAP-mcherry-FLAG-BirA* with the help of tissue-specific Gal4 driver lines were collected as described (Ramalingam et al., 2021). To isolate tracheal or gut cells from *lola* ^*-/-*^ embryos, fly lines containing *RAN-GAP-mcherry-FLAG-BirA* on the second chromosome (expressing in either trachea or gut) was recombined with the *lola-I* ^*ORC4*^ chromosome and maintained over a GFP-marked *CyO* balancer (Trachea: w[*]; P{w[+mC]=GAL4-btl.S}2, P{w[+m*]=lacZ-un8}276, p[UAS-3xFLAG-blrp-mCherry-RanGap, UAS-BirA)5; *lola*^orc4^/*CyO*-GFP) (Gut: w[*]; P{GawB}NP3084, p[UAS-3xFLAG-blrp-mCherry-RanGap, UAS-BirA)5; *lola*^orc4^/*CyO*-GFP). Homozygous embryos for the recombinant chromosome were obtained by sorting for GFP-negative embryos.

### Embryo collection and immunostainings

Adult fly maintenance and embryo collections were performed as described (Ramalingam et al., 2021). Briefly, embryos were collected and matured at 25 °C, then dechorionated for 1 min with 67% bleach and cross-linked for 15 min with 1.8% formaldehyde (final concentration in water phase). For the single-molecule FISH experiments, the embryos were cross-linked in 1x PBS in DEPC-treated water. Homozygous *lola-I* mutant embryos were obtained by sorting for GFP-negative embryos in PBT (PBS with 0.05-0.1% Triton). Embryos used for ChIP-seq experiments were also flash frozen in liquid nitrogen and stored at -80°C and used later. For ATAC-seq and scRNA-seq experiments, the embryos were processed immediately after dechorionation without crosslinking. For immunostainings, antibodies were used in the following dilutions: Lola-I (custom-made by Genescript) 1:750, α-tubulin antibodies (Sigma, T9026) at 1:500, Lamin (ADL40 from Developmental Studies Hybridoma Bank, DSHB, at 1:750), α-MHC (1:500), Elav (7E8A10 from DSHB) at 1:30, and Repo (8D12 from DSHB) at 1:10. For Western blot experiments, antibodies against Lola-I (custom-made by Genescript) were used 1:2000, those against Pol II (Rpb3, custom made from GeneScript, Zeitlinger lab 163185-50) at 1:2000.

### Isolation of tissue-specific nuclei

Nuclei isolation was performed using modified versions of previously published protocols (Bonn et al., 2012; Deal and Henikoff, 2011), as described in (Ramalingam et al., 2021).

### ChIP-seq experiments

Antibodies were raised against the Lola-I-specific portion (455-877 AA), thus excluding the BTB domain. It also excludes the DNA binding domains of Lola-I. Rabbit polyclonal antibodies against the full-length *Drosophila* Pol II subunit Rpb3 (custom made from GeneScript, Zeitlinger lab 163185-50) is also used. ChIP-seq experiments were performed as follows. ∼100 mg embryos were used per ChIP, and 5 μg chromatin was used for tissue-specific ChIP-seq experiments. Fixed embryos were homogenized by douncing in an ice cold A1 buffer (15 mM HEPES (pH 7.5), 15 mM NaCl, 60 mM KCl, 4 mM MgCl, 0.5% Triton X-100, 0.5mM DTT, protease inhibitors) and A2 buffer (15 mM HEPES (pH 7.5), 140 mM NaCl, 1mM EDTA, 0.5mM EGTA, 1% Triton X-100, 0.1% sodium deoxycholate, 0.1% SDS, 0.5% N-lauroylsarcosine, protease inhibitors) in a tissue grinder for 10-15 times in A1 and A2 buffer each. Then the sonication of the chromatin was performed with a Bioruptor Pico for four-five rounds of 30 seconds on and 30 seconds off cycles. The sonicated chromatin was cleared by centrifugation and the supernatant was used for ChIP. Chromatin was incubated with antibodies pre-bound to Dynal magnetic beads (IgA or IgG) overnight with end-to-end rotation at 4°C and washed with an ice-cold RIPA buffer (50 mM HEPES (pH 7.5), 1mM EDTA, 0.7% sodium deoxycholate, 1% NP-40 (IGEPAL CA-630), 0.5M LiCl). Eluted, reverse cross-linked DNA was then purified using phenol-chloroform-isoamylalcohol phase separation and ethanol precipitation. ChIP-seq libraries were prepared from 5-15 ng ChIP DNA or 100 ng input DNA according to the manufacturer ‘s instructions (NEBNext ChIP-Seq Library Prep kit).

### ATAC-seq and MNase-seq experiments

ATAC-seq was performed using ∼500-2000 embryos of stage 14–17 h AED. Nuclei were isolated by douncing the embryos in the HBS buffer as described above in the Isolation of tissue-specific nuclei section. Whole embryo ATAC-seq was performed without the selection of nuclei from a specific tissue using the *OregonR* embryos. The transposition of the nuclei was performed as described in (Buenrostro et al., 2013). Computational filtering for fragments of size 0-100 bp was done to capture signals from the accessible regions.

MNase digestion was performed similarly to previously published protocols (Mavrich et al., 2008). Briefly, chromatin was extracted from 0.1 mg of *Oregon-R* or *lola-I* mutant embryos per replicate by douncing embryos in the NPS buffer (0.5mM spermidine, 0.075% IGEPAL CA-630, 50mM NaCl, 10mM Tris-Cl (pH 7.5), 5mM MgCl, 1mM Cacl, 1mM beta-mercaptoethanol) using a tissue homogenizer, then digested with a concentration gradient of MNase (Worthington Biochemical Corporation #LS004798) in NPS buffer for 30 min at 37°C. All digestion concentrations were run on a gel and the concentration to be sequenced was chosen such that the digestion is complete, characterized by the presence of only mononucleosomes, but the samples are not over digested (no smaller than mononucleosome sized fragments). Libraries were prepared from purified MNAse digested DNA using the NEBNext DNA Library Prep kit following the manufacturer ‘s instructions and then paired-end sequenced on an Illumina HiSeq 2500 sequencing system. Computational filtering for fragments of size 100-200 bp to analyze the nucleosome occupancy.

### Nucleosome binding assay

Full-length Lola-I protein was expressed using baculovirus and purified by Genescript. Briefly, the *lola-I* sequence was synthesized and sub-cloned into the Flag-TAG expression vector F1 and expressed in Sf9 insect cells using a recombinant Bacmid. Sf9 cells were grown in Sf-900 II SFM Expression Medium in Erlenmeyer Flasks at 27°C in an orbital shaker. Cells were seeded in 6 wells, transfected the next day by adding DNA mixed with Cellfectin II at an optimal ratio, and incubated for 5-7 days before harvesting P1 and P2 viral stock. The Sf9 cells (1 L) containing 5% FBS were infected with the P2 virus at MOI=3 and harvested at 48 h post-infection. Cells were sonicated in 50 mM Tris, pH 8.0, 150 mM NaCl, 5% Glycerol containing protease inhibitors. Cell pellets were harvested and lysed, and the supernatant was incubated with Flag Columns to capture the target protein. Fractions were pooled and dialyzed with 50 mM Tris-HCl, 150 mM NaCl, 5% Glycerol, pH 8.0 followed by 0.22 um filter sterilization. Proteins were analyzed by SDS-PAGE and Western blot by using standard protocols for molecular weight and purity measurements.

The nucleosome binding assay was performed as previously described (Yu and Buck, 2019). Templates were designed by altering the right side of the Widom 601 nucleosome positioning sequence and placing Lola-I binding motifs with increasing distance to the dyad axis (Supplementary Table 1). Four translational settings were tested – dyad (at superhelix location (SHL) R0, R0.5), intermediate (SHL R4, R4.5), edge (SHL R6, R6.5, R7), and linker, which is outside the nucleosome (SHL 8). The rotational setting of each motif was designed such that it is either outside on the solvent accessible side (SHL R0.5, R4.5, R6.5) or concealed (SHL R0, R4, R6, R7) based on the nucleosome crystal structure formed on the Widom 601 sequence (Makde et al., 2010). To explore cooperativity, a template with two neighboring motifs was designed (SHL R6+R7), as well as one with two motifs further apart as a control (SHL L6 on left +R7). All templates were compared to non-specific binding to the Widom 601 sequence.

All 11 synthesized DNA templates were amplified via PCR with the primer pair 5 ‘-GATGGACCCTATACGCGGC-3 ‘ and 5 ‘- GGAACACTATCCGACTGGCA-3 ‘, and the products were column-purified (QIAGEN), quantified, and pooled equally. *In vitro* nucleosomes were generated from H2A/H2B dimer and H3.1/H4 tetramer (NEB). The pool of 11 nucleosome sequences were added to histones at octamer/DNA molar ratios of 1.25:1 in 2M NaCl solution. Nucleosomes were reconstituted by salt gradient dialysis as described in (Hayes and Lee, 1997), purified from free DNA with 7%- 20% sucrose gradient centrifuge, and concentrated by 50K centrifugal filter units (Millipore, AmiconR Ultra).

For the protein-nucleosome binding assays, each of 0.25 pmol of purified nucleosomes were incubated with increasing concentrations of Lola-I protein (molar ratios of 0:1, 1:1, 2:1, 4:1 to 10:1) in 7 μl DNA binding buffer (10mM Tris-Cl, pH7.5; 50mM NaCl; 1mM DTT; 0.25mg/ml BSA; 2mM MgCl2; 0.025% P-40; and 5% glycerol) for 10 minutes on ice and then for 30 minutes at room temperature. Protein binding was detected by mobility shift assay on 4% (w/v) native polyacrylamide gels (acrylamide/bisacrylamide, 29:1, w/w, 7 × 10 cm) in 0.5 × Tris Borate-EDTA buffers at 100V at 4 °C. After electrophoresis, DNA was imaged by staining with SYBR Green (LONZA).

All visual bands were excised from the gel, as well as the bands at the same locations in the other lanes. Each gel slice was processed separately for a total of 20 samples from 2 replicate experiments. In order to extract DNA from polyacrylamide gel, the chopped gel slices were soaked in diffusion buffer (0.5 M ammonium acetate; 10mM magnesium acetate; 1mM EDTA, pH8.0; 0.1% SDS), and incubated at 50 °C overnight. The supernatant was collected, residual polyacrylamide removed with glass wool, and DNA purified with QIAquick Spin column (QIAGEN). The DNA concentration for each sample was determined by qPCR by comparing it to a standard curve generated from the control 601 sequence. Based on this concentration, the samples were amplified by PCR using Illumina primers (the cycle number ranged from 8 to 12) and then indexed in a second round of PCR using Nextera dual index primers, followed by clean up with AMPure XP beads (Beckman Coulter). The samples were multiplexed and sequenced on an Illumina MISeq using 2 × 150-bp paired-end sequencing. Sequencing and quality control were performed at the University at Buffalo Genomics and Bioinformatics Core.

High-quality sequence reads were mapped to each specific starting sequence using VSearch (Rognes et al. 2016). After obtaining the amount of the reads from each band, the data from each of the nucleosome sequences N were normalized to the Widom 601 control sequences: The results were then analyzed relative to the 601-control sequence and the non-specific binding in the input lane without any Lola-I. Relative shift is determined from the non-shifted nucleosome bands and controls for the technical variability introduced by gel-excision, PCR, NGS-library construction, or NGS sequencing. In this method each specific nucleosome sequence or the super-shifted sequence is measured relative to non-specific binding (601 fragment without a Lola-I binding site):

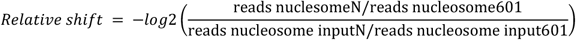

where N is one of the 11 nucleosome sequences, *601* is the control nucleosome sequence, *reads nucleosome* is the nucleosome band at a specific concentration of Lola-I, *reads nucleosome input* is the nucleosome band in the input lane without any Lola-I added. Or as,

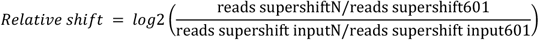

where N is one of the 11 super-shifted sequences, *601* is the control nucleosome sequence, *reads supershift* is the *super-shift*ed band at a specific concentration of Lola-I, *reads supershift input* is the *super-shift*ed band in the input lane without any Lola-I added.

### Single-molecule FISH experiments

Stellaris single-molecule FISH probes were designed for the *Drosophila melanogaster* genes *Gip, PPO1*, and *PPO2* using the Stellaris probe designer, and purchased with a label ready C-term TEG-Amino tag from Biosearch Technologies. 4 nMol of *PPO1* and *PPO2* probe sets were combined and labeled with two units of AF647 amine reactive succinimidyl ester Decapacks (ThermoFisher) and the *Gip* probe set was labeled similarly, with AF555 and *lola-I* with AF647. Labeling was overnight at 4 °C in 0.1 M sodium tetraborate at pH 9. HPLC was used to purify labeled from unlabelled probes as described (De Kumar et al., 2017).

The single-molecule FISH technique was optimized and adapted from the Stellaris website and based on Dr. Shawn Little ‘s protocol (personal communication). Embryos were collected, dechorionated with bleach and crosslinked in 4% formaldehyde in a 1:3 mixture of PBS and heptane. Embryos were then sorted in PBT (0.1% Triton), and then devitellinated with a 1:1 mixture of methanol and heptane, equilibrated in methanol and stored at -20 °C. After gradual rehydration, embryos were post-fixed, treated with proteinase K (0.5ug/ml) for 1 h on ice and 1 h at 37 °C, and crosslinked once more. The permeabilized embryos were washed with a series of PBT and Stellaris WashA buffers. The embryos then underwent prehybridization in the Stellaris Hyb buffer for one day at 37 °C. Hybridization with 1 μM concentration of probes was performed at 25 °C overnight. After several washes with WashA buffer and one with PBS, DAPI staining was performed in PBS buffer (5 μg/ml) for 5 minutes, followed by washes with PBT buffer to remove unbound DAPI. Embryos were mounted in Prolong gold and cured at room temperature (22 °C) for three days to a week.

For analyzing *lola-I* transcripts, images of whole-mount *Drosophila* embryos were acquired on a PerkinElmer Ultraview spinning disk confocal microscope equipped with an EM-CCD camera (model C9100-13; Hamamatsu Photonics), using Volocity software (PerkinElmer). An Apochromat 63x, 1.46 NA oil immersion objective was used with a 405/488/561/640 nm multiband dichroic. Dual color images of DAPI and *lola-I*-Af647 were acquired with 405 nm and 640 nm laser lines, respectively, with 415-475 nm emission filter for Dapi and a 660-750 nm emission filter for Af647.

For the expression analysis, images of whole-mount *Drosophila* embryos were acquired from a Nikon 3PO spinning disc, with a W1 disc, sitting on a Nikon Ti Eclipse base, controlled by Nikon Elements. Data were collected either with a 40x 1.1 NA Plan ApoChromatic long working distance water objective (for overviews), or a 100X, 1.4 NA Plan-Apochromatic oil objective (for observation of nascent transcripts, single transcripts, and for transcriptional modeling). AF647, AF555, and DAPI were excited with 640 nm, 561 nm, and 405 nm lasers, respectively, through a 405/488/561/640-nm main dichroic. Emission filters included a 700/75- and 455/50-nm dual band pass filter for the far red and DAPI channel, and a 605/70 nm emission filter for the red channel. An ORCA-Flash 4.0 V2 digital sCMOS camera was used, Z-step spacing was 1 micron for the 40x overview data, and 0.3 microns for 100x data. For 40x data, prior to display, gut autofluorescence was subtracted from the red channel using a reference in the green channel.

For quantification of transcripts per cell for transcriptional modeling, the 100x data were used. A Gaussian blur of radius 1 pixel was applied, followed by a rolling ball background subtraction with a radius of 50 pixels. To integrate the total signal over each cell, a z bin of 7 was applied for a final spacing of 2.1 microns. Cell outlines were manually drawn in FIJI, and integrated intensity was taken over 3 of the binned z slices, for a total cell size in z of 6.3 microns, after an intensity threshold was applied to remove the background. This background intensity was found from areas with no visible transcripts. Total cells counted were 184 from 7 wild-type embryos, and 340 cells from 14 *lola-I* embryos of 12-14h. To calibrate transcripts per cell from integrated intensity, single transcripts were fit to a 2-dimensional Gaussian. The total integrated intensity per cell was then divided by the average integrated intensity of single transcripts to get the number of transcripts per cell. Likewise, to find the intensity of nascent sites, nascents were identified and fit to a 2-dimensional Gaussian. The integrated intensity of the Gaussian was then divided by a single RNA spot intensity to yield the number of RNAs per nascent.

The fit of the distributions of transcripts per cell to the simple 2-state model shown in Figure S3C was done as described (Raj et al., 2006):

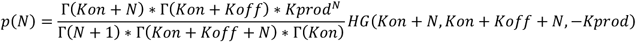

where, *HG* is the confluent hypergeometric function of the first kind, *K*_*on*_ is the activation rate constant, *K*_*off*_ is the inactivation rate constant, and *K*_*prod*_ is the production rate constant, all expressed as ratios to the degradation rate constant. Given the difficulty in fitting these complex distributions, we took advantage of the fact that we can observe nascent transcripts for transcriptionally active cells. That lets us measure the ratio of active to inactive cells and therefore the ratio of activation to inactivation rates:

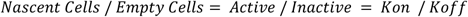

where, the ratio is fixed in the analysis to calculate K_off_ from K_on_. Fitting was accomplished with the scipy optimize curve_fit function in python utilizing the trf (trust region reflective) algorithm (Branch et al., 1999; Moré, 1978). Because the variance is higher for more frequent histogram bins and those are the bins that are most confidently determined for our data set, fitting was accomplished without weighting. Errors are determined by Monte Carlo analysis. This analysis assumes that the data distribution is reasonably well described by the fit and simulates random distributions repeatedly from the fit distribution and fits those distributions to obtain the error distribution for the fit parameters. For wild-type data, the fit produced the following parameters relative to the degradation rate constant: an activation rate constant, K_on_, of 2.7 +/- 0.42 and a production rate constant, K_prod_, of 1524 +/- 58. From the images, we know the ratio of active/inactive is 0.69. Taking these data together yields an inactivation rate constant, K_off_, of 3.7. For the mutant, the fit produced a K_on_ of 0.19 +/- 0.12 and a K_prod_ of 571 +/- 207.8. From the images we know that the ratio of active/inactive is 0.036. These values yield a K_off_ of 5.1. Supplemental table 3 shows the values of these parameters and expected values based on reasonable literature mRNA half-lives.

### mRNA-seq and scRNA-seq experiments

mRNA-seq and scRNA-seq experiments were performed as described (Karaiskos et al., 2017; Ramalingam et al., 2021) with isolated RNA from entire embryo or single cells from 14-14.5h AED *Oregon-R* embryos or *lola-I* mutant embryos sorted for homozygous mutant embryos that are GFP^-^. Total mRNA was extracted from non-cross-linked embryos using the Maxwell Total mRNA purification kit (Promega, #AS1225). mRNA from single cells for the scRNA-seq experiments was isolated using the 10x Genomics instrument.

### Sequence alignment

Reads from the ChIP-seq data and ATAC-seq experiments were aligned to the Drosophila melanogaster genome (dm6) using Bowtie (v 1.1.2) (Langmead et al., 2009), allowing a maximum of two mismatches and including only uniquely aligned reads. Coverage files were created by extending the aligned reads to the estimated insert size or the actual size for the paired-end libraries. For the bulk mRNAseq samples, pseudo-alignment was performed using the Kallisto package (Bray et al., 2016), to calculate the gene expression values. For the scRNA-seq samples, alignment and separations of reads from different cells and quantification of gene expression were done using the Cell Ranger pipeline (v 2.1.1) from 10x Genomics.

### Analysis of single-cell RNA-seq data

The wild-type scRNAseq data were aligned to the *lola* ^*-/-*^ scRNAseq data by performing the canonical correlation analysis (CCA) using the Seurat package. Gene expression in a particular tissue/ single-cell cluster is measured either using the median absolute normalized expression counts or using the percentage of cells with any detectable transcripts for each gene in each tissue. Expression of each Lola-I target gene was calculated in the presumed target tissue (sc-cluster with the maximum expression for each gene was considered as the expressing tissue for that gene) and the presumed non-expressing tissues (the five least expressing sc-clusters for each gene were considered as other tissues for that gene).

### Gene groups

Control genes are a randomly chosen 250 gene-subset of the constant set genes (Gaertner et al., 2012). Lola-I binding sites are divided into promoter proximal and distal sites based on whether the binding sites are present within +50 to -450 bp of an annotated TSS (flybase version r6.21). Lola-I targets are defined as genes with a Lola-I binding peak (using the MACS2 peak caller) in both the Lola-I ChIP-seq replicates in the promoter region. Lola-I targets genes are further filtered for at least a two-fold change in pol II occupancy between wild-type and mutant embryos for the scRNAseq analysis.

### Motif enrichment analysis

*De-novo* identification of the Lola-I motif (position weight matrix) found at the opening-set genes in figure 1B was done using the MEME package (meme macs_peaks_late_genes.fasta -oc macs_peaks_late_genes_meme_output -p 5 -mod zoops -dna -nmotifs 10 -revcomp -maxw 12 -maxsize 5000000). Enrichment analysis of the Lola-I motif was done by scanning for the AAAGCTY motif. Unbiased enrichment analysis of all known motifs from the MotifDb package were scanned using the FIMO package and then the enrichments for each of the motifs was tested using a chi-squared test.

### Statistical significance calculations and data visualization

*P* values in figures 1D, 1G, 2D, 2E, 4A, 4B, 4C, S1B, S1C, S1D, S1G, S4A were calculated with the two-sided Wilcoxon test. *P* values in figure 1A were calculated with the chi-squared test with multiple-testing correction. *P* values in Supplementary Table 2 were calculated using the hypergeometric test. Heat maps are normalized, and really low or high values are ceiled or floored, respectively. Box plots show the median as the central line, the first and the third quartiles as the box, and the upper and lower whiskers extend from the quartile box to the largest/smallest value within 1.5 times of the interquartile range.

### Data and software availability

Raw and processed data associated with this manuscript have been deposited in GEO under session number GSE200875 and will be available after peer-reviewed publication. All data analysis performed in this paper, including raw data, processed data, software tools, and analysis scripts will also be available through publicly accessible Amazon Linux virtual machine image after peer-reviewed publication. The analysis code is also available on GitHub at https://github.com/zeitlingerlab/Ramalingam_Lola_2022.git

## Acknowledgments

We thank Cindi Staber for the help in designing and coordinating the Lola-I protein and Lola-I antibody production with Genescript. We thank Shawn Little for sharing his single-molecule FISH protocol. We thank Kate Hall, Allison Peak, and Ana Pinson for the technical assistance with the scRNA-seq experiments. We thank Divya Krishna Kumar for the technical assistance with the single-molecule FISH experiments. We thank Robb Krumlauf and Mounia Lagha for their feedback on the manuscript. This work was supported by funding from the National Institutes of Health (DP2OD004561) and the Stowers Institute for Medical Research. This work been done to fulfill, in part, requirements for Vivekanandan Ramalingam ‘s PhD thesis research as a student of the University of Kansas.

## Author contributions

V.R. and J.Z. conceived the project, designed the study and wrote the manuscript. V.R. performed all the experiments and the analysis under the supervision of J.Z., except for the nucleosome binding assays. X.Y., under the supervision of M.B., performed the nucleosome binding assays, including the library preparation and the analysis of the data. B.S., J.U. and J.J.L. helped with the single-molecule FISH experiments, the image acquisition and performed the analysis of the data, K.J.B. analyzed the phenotype of the *lola-I* mutant embryos using the immunostaining assay, A.O. helped to develop the single-molecule FISH protocol, M.N. helped with the tissue-specific ChIP-seq experiments. All authors provided input on the manuscript.

## Supplementary figures

**Figure S1:**
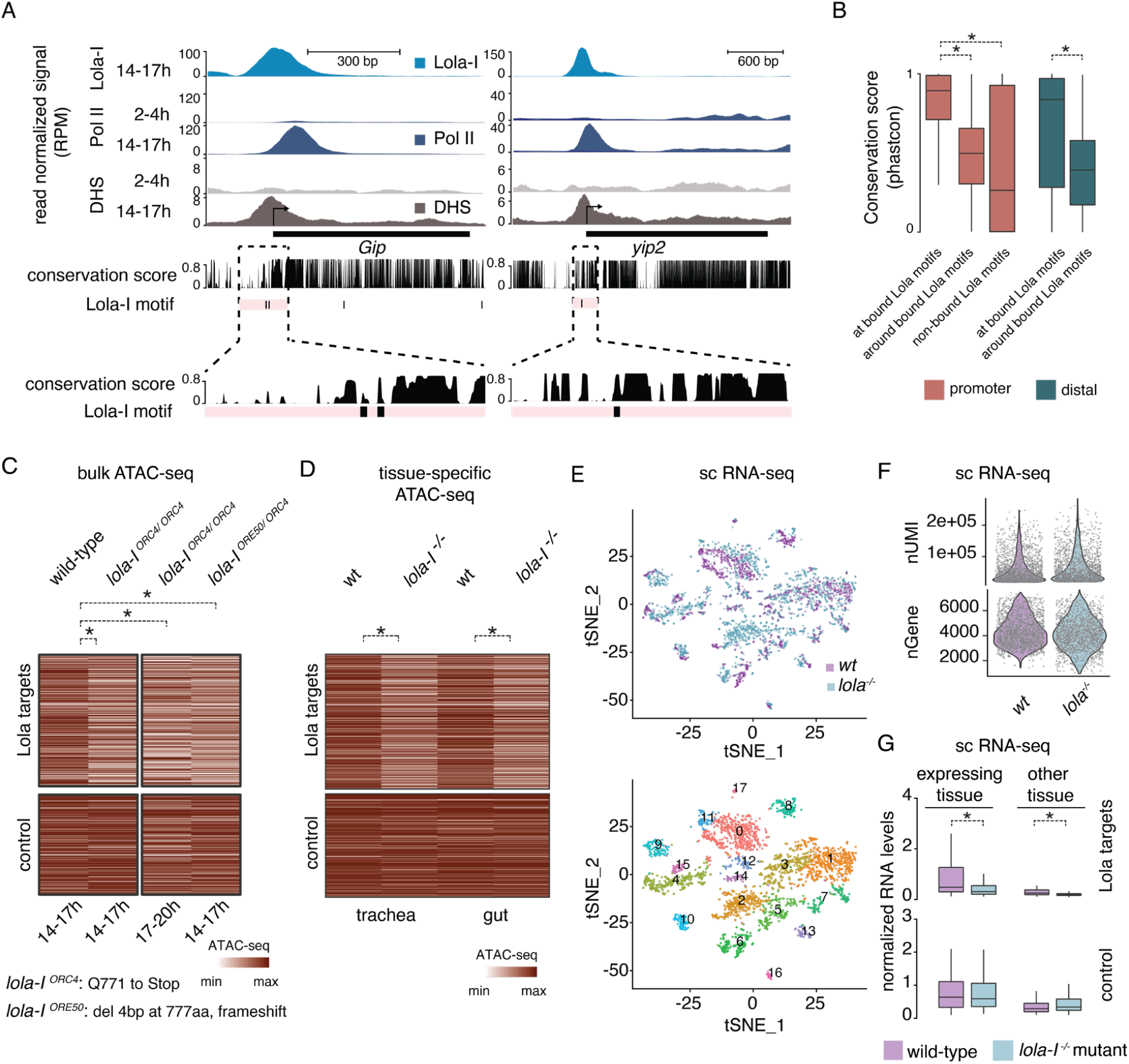
Conservation of Lola-I motifs and genomic characterization of *lola-I* mutants. **A)** Single-gene examples of Lola-I target promoters showing Lola-I binding, Pol II binding, DNase hypersensitivity (DHS) and the conservation of Lola-I binding sites across different *Drosophila* species using the PhastCons score. **B**) Boxplots of PhastCons scores at Lola-I motifs show significant conservation of bound Lola-I motifs. Bound Lola-I motifs at promoters are highly conserved compared to promoter regions in general (100 bp centered on the motif) or non-bound Lola-I motifs in promoters. A Wilcoxon two-sided test was used and the star (*) denotes P <10^−15^. Similarly, Lola-I binding sites at distal (non-promoter) regions bound by Lola-I are also highly conserved (Wilcoxon two-sided test, *P <10^−15^). **C)** Heatmap showing ATAC-seq accessibility at Lola-I targets and control promoters for wild-type and mutant combinations. Lola-I targets show reduced accessibility in *lola*^ORC4^ homozygous mutant embryos over wild-type at 14-17h (Wilcoxon two-sided test, *P <10^−9^) and this extends to 17-20h (Wilcoxon two-sided test, *P <10^−10^), confirming that the opening of these promoters is not just delayed in *lola-I* mutants. Importantly, trans-heterozygous *lola* ^ORE50/ORC4^ mutant embryos (14-17h) show a similar reduction in accessibility compared to homozygous *lola* ^ORC4/ORC4^ embryos (Wilcoxon two-sided test, *P<10^−13^), demonstrating that the reduced accessibility is due to the mutation in *lola-I* and not due to other mutations in the *lola* ^ORC4^ line. *lola* ^ORC4^ mutant has a premature stop codon before the DNA binding domain. *lola*^ORE50^ mutant has a frameshift mutation before the DNA binding domain. **D)** Heatmap showing reduced ATAC-seq accessibility at Lola-I targets in *lola* ^-/-^ mutant embryos over wild-type at 14-17h when tracheal nuclei (left) or gut nuclei (right) were isolated using the INTACT method (Wilcoxon two-sided test, *P<10^−9^) These tissues were selected because the driver lines expressing nuclear-envelope BirA in these tissues are located on the second chromosome and thus could be recombined with the *lola-I* mutants. The results show that the reduced ATAC-seq accessibility of Lola-I target promoters in *lola-I* mutants is not tissue-specific. **E)** Single-cell RNA-seq (scRNA-seq) tSNE map of cells from wild-type and *lola-I* mutant homozygous embryos are shown. Top panel: the projection of the scRNA-seq data from both replicates shows alignment between the wild-type and *lola-I* mutant cells. Bottom panel: the identified clusters in these scRNA-seq data show that wild-type and *lola-I* mutant cells cluster together. **F**) Number of Unique Molecular identifiers (UMI) and number of genes captured per cell are comparable between the wild-type and *lola-I* mutant samples, indicating similar data quality. **G**) Boxplots of scRNA-seq data show that both the tissue-specific expression and the basal expression of Lola-I targets are reduced in the *lola-I* embryos compared to wild-type embryos (both profiled at 14-14.5h (Wilcoxon two-sided test, expressing tissue - *P<0.02, other tissues - *P<0.003). Normalized expression in each cell shown. Box plots in B and G show the median as the central line, the first and the third quartiles as the box, and the upper and lower whiskers extend from the quartile box to the largest/smallest value within 1.5 times of the interquartile range.

**Figure S2:**
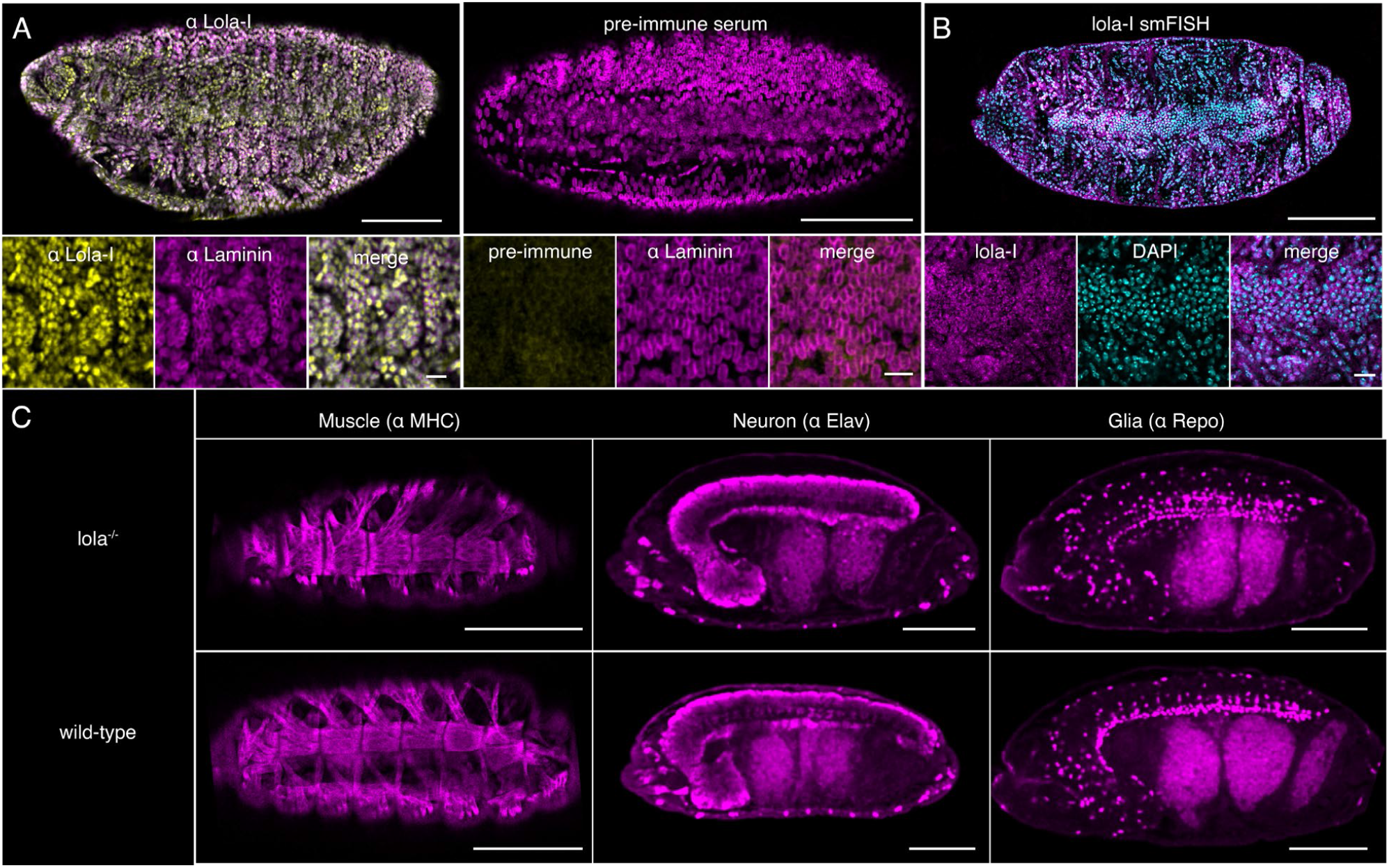
Immunostainings showing ubiquitous Lola-I expression and a *lola-I* mutant phenotype that appears anatomically normal. Immunostainings using antibodies against Lola-I (1:750) and Lamin (1:750) as control show ubiquitous nuclear signals of Lola-I throughout the embryo (left). Control Immunostainings with the pre-immune serum from the Lola-I antibody production shows no nuclear signal (right). scale bar: top - 100 μm and bottom - 10 μm **B**) Single-molecule FISH also shows ubiquitous expression of *lola-I* transcripts. scale bar: top - 100 μm and bottom - 10 μm). **C**) *lola-I* mutants do not show any gross visible phenotypic defects but fail to hatch. Embryos were immunostained to visualize muscle (alpha-MHC antibodies), neurons (Elav antibodies) and glia (Repo antibodies), respectively. scale bar: 100 μm. For all images here the contrast was adjusted linearly in every image and set the same in panel A (overviews and zooms) for **α** Lola-I and pre-immune serum and separately for **α** Laminin. Additionally, the lookup tables for all images are linear. The settings in the individual panels are the same as in the merge.

**Figure S3:**
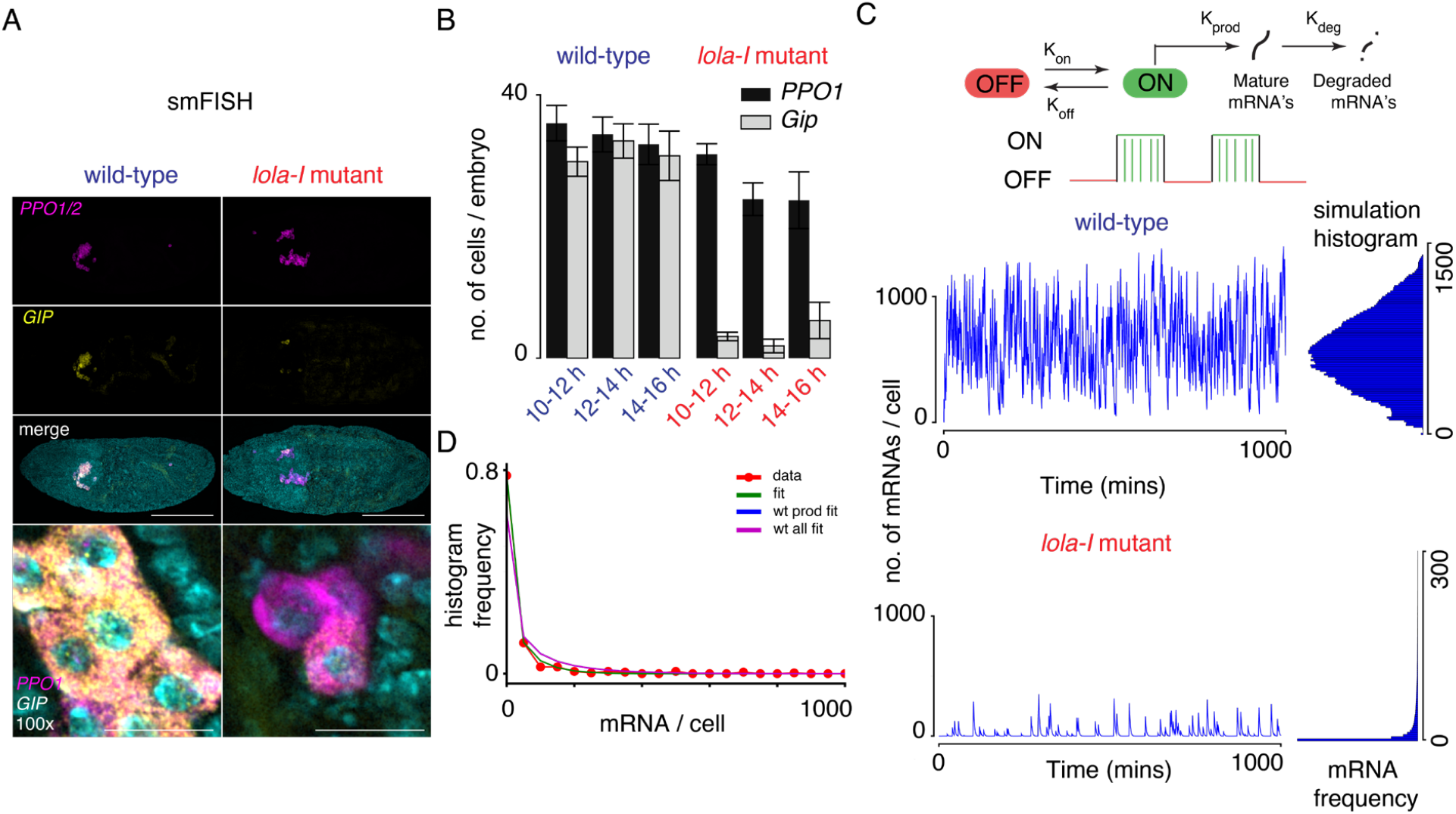
Quantification of expression changes in Lola-I mutants using single-molecule FISH. **A**) Example of single-molecule FISH images of *Gip* (yellow), and *PPO1* and *PPO2* genes (pink) in wild-type (wt) embryos and *lola-I* mutant embryos. Data were acquired with 10x magnification (top three panels); scale bar = 100 μm and 100x magnification (bottom panel); scale bar = 20 μm. Here the images were all brightness and contrast adjusted for clarity and linearly. The lookup tables for each image are linear. The settings in the individual panels are the same as in the merge. **B)**The number *Gip-*positive cells are strongly reduced in *lola-I* mutant embryos compared to wild-type, while the *PPO1*/*PPO2* positive cells remain similar. Bar plots show data for wild-type: 10-12 h n=7, 12-14 h n=8, 14-16 h n=4, *lola-I* mutant: 10-12 h n=7, 12-14 h n=7, 14-16 h n=7. Average is shown as a bar with standard error of the mean as error bars. Reduced *Gip* expression is observed for multiple time-points showing that it is not due to a developmental delay and that *Gip* expression only mildly recovers over time. **C**) Simulations of bursty transcription for a single cell (left) and the histogram of mRNA levels (right) for wt fit parameters (top row) and mutant fit parameters (bottom row). The model of transcription used in the simulations and fits of experimental histograms. Gene activation (with rate constant, K_on_) leads to the initiation of a mRNA production burst with the appearance of nascent spots. Gene inactivation (with a rate constant, K_off_) causes the disappearance of nascent spots and leads to the decay of mature mRNA levels (due to degradation) at the end of a burst. Wild-type bursts are so frequent that one rarely finds dark cells, while mutant bursts are infrequent, leading to mostly dark cells. D) Distribution of the numbers of mRNAs per cell quantified from the total fluorescence intensity (divided by the intensity of a single RNA molecule) of the single-molecule FISH signal from each cell in the *lola-I* mutant embryos (yellow dots). Here, many *PPO1/PPO2* positive cells were dark for *Gip* signal. The distribution was fit according to the two-state model (see methods). In one case the data was fit with all parameters allowed to converge (green line). In another case the data was fit with the production rate fixed to wild-type while all other parameters were allowed to converge (blue line). In the final case, all all adjustable parameters were fixed to the wild-type values (purple line).

**Figure S4:**
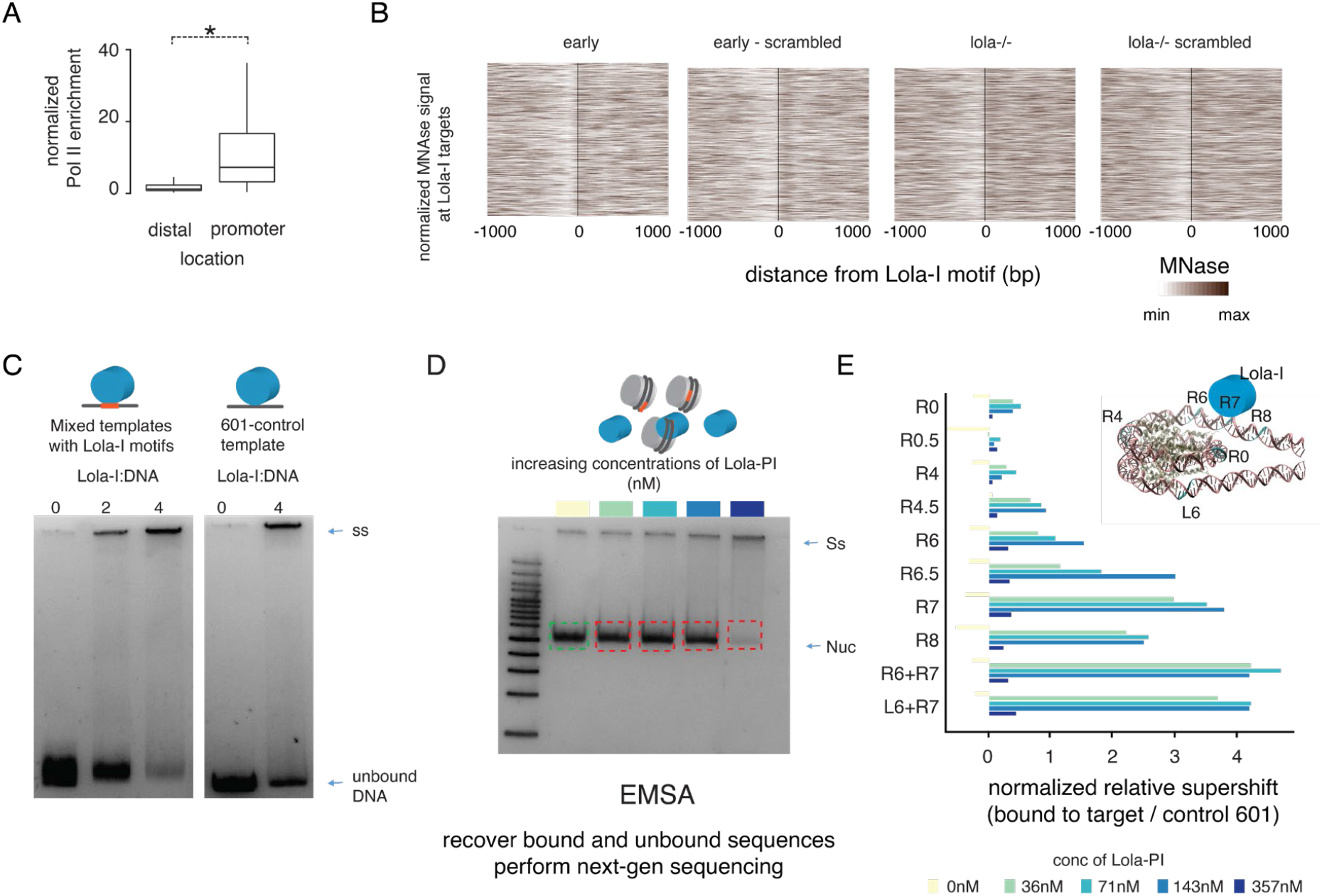
Additional information on in vitro nucleosome binding assays. **A**) Box plot shows normalized Pol II enrichment (wt 14-17 h) at both the promoter proximal and distal Lola-I peak regions. Pol II signal is found only at the annotated promoter regions and not at the distal Lola-I target regions (Wilcoxon two-sided test, *P <10-15). Box plot shows the median as the central line, the first and the third quartiles as the box, and the upper and lower whiskers extend from the quartile box to the largest/smallest value within 1.5 times of the interquartile range. **B**) Nucleosome occupancy centered on Lola-I motifs or randomly scrambled (−85 or +85 bp) are plotted as MNase-seq heatmap at each Lola-I binding site for the early (2-4 h) wt or late (14-17 h) Lola-I mutant samples. The left-right orientation is chosen such that the nucleosomes on either side of the motif align. The results suggest that the Lola-I motifs might be preferentially found along the nucleosomal edges in vivo. **C**) Binding of Lola-I to the 601-templates with Lola-I motif (left panel) or 601-control template without any Lola-I motif (left panel) at different concentrations of Lola-I is measured. Lola-I binds to the 601-templates with the Lola-I motifs at lower concentration than to the 601-control template. **D**) The gel-shift assay shows the binding of Lola-I to nucleosome bound 601 sequences with differing Lola-I motifs. Both the nucleosomal and the super-shifted fractions are purified and sequenced to measure the relative affinity of Lola-I to different Lola-601 sequences. **E**) Relative super-shift for Lola-I binding at different concentrations to nucleosomes with 601-Lola-I sequences with Lola-I motif vs control-601 template without the Lola-I motif (relative to the no Lola-I lane), are shown. Lola-I strongly binds when the motif is located along the nucleosomal edge. At the highest Lola-I concentration, non-specific binding occurs, which reduces the relative super-shift values.

